# The rust fungus *Melampsora larici-populina* expresses a conserved genetic program and distinct sets of secreted protein genes during infection of its two host plants, larch and poplar

**DOI:** 10.1101/229971

**Authors:** Cécile Lorrain, Clémence Marchal, Stéphane Hacquard, Christine Delaruelle, Jérémy Pétrowski, Benjamin Petre, Arnaud Hecker, Pascal Frey, Sébastien Duplessis

## Abstract

Mechanims required for broad spectrum or specific host colonization of plant parasites are poorly understood. As a perfect illustration, heteroecious rust fungi require two alternate host plants to complete their life cycle. *Melampsora larici-populina* infects two taxonomically unrelated plants, larch on which sexual reproduction is achieved and poplar on which clonal multiplication occurs leading to severe epidemics in plantations. High-depth RNA sequencing was applied to three key developmental stages of *M. larici-populina* infection on larch: basidia, pycnia and aecia. Comparative transcriptomics of infection on poplar and larch hosts was performed using available expression data. Secreted protein was the only significantly over-represented category among differentially expressed *M. larici-populina* genes in basidia, pycnia and aecia compared together, highlighting their probable involvement in the infection process. Comparison of fungal transcriptomes in larch and poplar revealed a majority of rust genes commonly expressed on the two hosts and a fraction exhibiting a host-specific expression. More particularly, gene families encoding small secreted proteins presented striking expression profiles that highlight probable candidate effectors specialized on each host. Our results bring valuable new information about the biological cycle of rust fungi and identify genes that may contribute to host specificity.

## Introduction

Rust fungi (Pucciniales) are obligate biotrophs that establish parasitic associations with their host plants. Pucciniales depict one of the largest order of plant pathogenic fungi with more than 8,000 species infecting a wide range of hosts within ferns, gymnosperms and angiosperms (Aime et al., 2014; Cummins and Hiratsuka, 2003). A remarkable feature of some rust fungi is heteroecism – they infect two unrelated host plants to complete their life cycle. To do so, they produce different spore forms and the life cycle is qualified of macrocyclic. For instance, the poplar rust fungus *Melampsora larici-populina* has a heteroecious and macrocyclic life cycle (Hacquard et al 2011a; Vialle et al., 2011). It infects two unrelated host plants: Larch (*Larix* spp., conifer, aecial host) and Poplar (*Populus* spp., dicot, telial host). Larch infection is realised by *M. larici-populina* haploid basidiospores and results in the formation of pycnia and pycniospores (Pinon & Frey, 2005; Hacquard et al. 2011a). After fertilization and production of aecia, airbone aecispores infect poplar leaves leading to the production of urediniospores within a week (Harder, 1984; Voegele et al., 2009). Asexual multiplication of urediniospores causes severe epidemics in poplar plantations all over summer (Pinon & Frey, 2005). In autumn, uredinia differentiate into telia, in which karyogamy occurs and diploid teliospores overwinter on dead poplar leaves (Hacquard et al., 2013). When favourable conditions are met the next spring, basidiospores are produced and the cycle repeats again. The molecular bases of these tightly controlled developmental stages and the genetic programmes underlying host alternation in rust fungi remain poorly understood (Bakkeren et al., 2016; Liu et al., 2015; Xu et al., 2011).

During infection, pathogens use effector proteins to modulate the cellular and the immune processes of the host plant (Win et al., 2012; Lo Presti et al., 2015). Effectors are critical to ensure successful establishment of the pathogen in host tissues (Lo Presti et al., 2015). In filamentous plant pathogens (fungi and oomycetes), effector proteins are often small proteins that possess a signal peptide for secretion (Sperschneider et al., 2015). In rust fungi, genome and transcriptome analyses predicted hundreds of secreted proteins (SPs) representing candidate effectors expressed during host infection (Duplessis et al., 2012; Duplessis et al., 2014). Effectors are specifically adapted to manipulate host plants and they evolve through the pressure of host recognition receptors (Rovenich et al. 2014).

The genetic programme used by *M. larici-populina* to colonize poplar leaves has been extensively described (Joly et al., 2010; Duplessis et al., 2011b; Hacquard et al., 2010, 2012; Petre et al., 2012); mostly because of epidemics occurring in poplar plantations (Pinon and Frey, 2005). Transcriptome analyses revealed waves of expression for candidate effector genes at different time-points on poplar, suggesting that the fungus tightly controls its genetic program during infection (Duplessis et al., 2011b; Hacquard et al., 2012; Petre et al., 2012). These studies helped to prioritize candidate effectors for functional analysis (Petre et al., 2015a; Petre et al., 2015b; Germain et al., 2017). However, it is not known whether similar or different sets of effectors are acting during the infection of the aecial and the telial host of *M. larici-populina*. This has been a general and recurrent question related to the capacity of heteroecious rust fungi to infect different hosts belonging to unrelated plant taxa (Schulze-Lefert and Panstruga, 2011; Duplessis et al., 2014). So far, a limited number of studies addressed gene expression in the different hosts of rust fungi (Cuomo et al., 2017; Liu et al., 2015; Xu et al., 2011) and none clearly determined to which extent a portion of the gene complement may be unique to one host or another.

In this study, we took advantage of the recent ability to complete the life cycle of the poplar rust fungus under laboratory conditions (Pernaci et al., 2014) to investigate the genetic programme used by the fungus during larch colonization. To this end, we used an Illumina RNA-sequencing (RNAseq) approach to assess fungal transcript abundance at three different stages of the life cycle: basidia, pycnia, and aecia. In a second time, we took advantage of previously published transcriptome data during poplar infection to compare *M. larici-populina* gene expression profiles on the two host plants. We identified genes preferentially expressed on larch or poplar, including secreted proteins that represent potential specific candidate effectors. We further scrutinized the expression features of pre-defined Small-Secreted Protein (SSP) gene families (Hacquard et al., 2012) to investigate whether members from SSP families showed a host-specific expression.

## Material & Methods

### Experimental setup and inoculation procedures

Specific *M. larici-populina* isolate 98AG31 stages were collected on the aecial host larch (*Larix decidua*) following the procedure reported in Pernaci et al. (2014). The Figure 1 describes the experimental set up and the Figure S1 details the overall procedure used to produce the biological material. Basidia were collected from dead poplar leaves and contained remains of telia and freshly produced basidia and basidiospores. Pycnia were collected from larch needles seven days after inoculation with basidiospores and are made of different fungal cell types such as *in planta* infection hyphae, flexuous hyphae and pycniospores. Aecia were collected from larch needles two weeks after inoculation and corresponded to freshly produced aecia and remaining pycnia. Three biological replicates were harvested at each stage; basidia replicates came from different poplar leaves, while pycnia and aecia replicates were produced from different boxes of inoculated larch seedlings. For each replicate (i.e. R1, R2, R3), aecia and pycnia were harvested from different larch seedlings present in the same inoculated box, ensuring synchronization of these stages after a single initial basidiospore inoculation.

**Figure 1:**
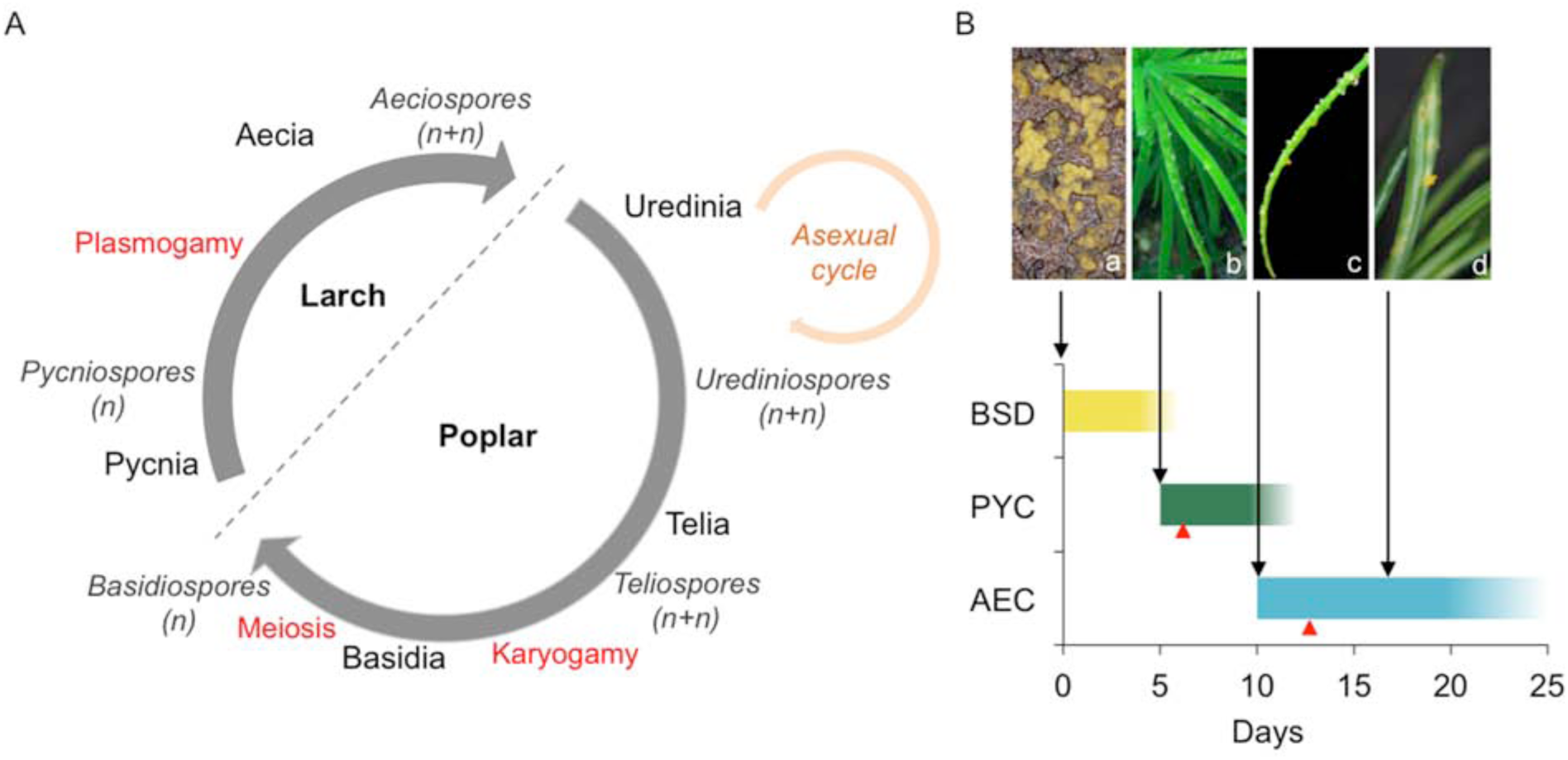
*Melampsora larici-populina* life cycle and experimental design for stage selection. (A) *M. larici-populina* heteroecious and macrocyclic life cycle. Haploid basidiospores (n) infect aecial host (*Larix spp*), (0) Pycnia are formed in larch needles producing nectar droplets containing pycniospores (n) with different mating types for fertilization. After plasmogamy, dikaryotic aecia establish. Aeciospores (n+n) infect leaves of the telial host (*Populus spp*) and produce dikaryotic urediniospores (n+n). Asexual cycle with continuous clonal production of urediniospores takes place during the summer season. In autumn, telia form teliospores (n+n) where karyogamy occurs and meiosis begins before dormancy. After winter, meiosis continues, basidia germinate and produce haploid basidiospores (n). (B) Pictures from left to right: (a) basidia and basidiospores, (b) pycnia nectar droplets, (c) aecia and aeciospores and (d) a few remaining pycnia, aecia and aeciospores. Color bars indicate the three larch-infection related stages sampled for RNA sequencing. BSD refers to basidia and basidiospores and the yellow bar represents the time between basidiospores inoculation and pycnia formation. PYC corresponds to pycnia and pycniospores (visible as nectar droplets) and the green bar represents the period of presence of visible PYC on larch needles. AEC refers to aecia and aeciospores and the blue bar indicates the presence of AEC on larch needles. Red arrows represent the time of collection for the two larch-infection stages PYC and AEC.

### RNA isolation and Illumina RNA-Sequencing

Total RNA from basidia samples were isolated with the RNeasy Plant Minikit (QIAGEN, Courtaboeuf, France) using RLC buffer with β-mercaptoethanol following manufacturer’s recommendations. For each basidia replicate, several rounds of RNA isolation were performed using low inputs of material (10 to 20mg) to avoid RNA degradation from decayed leaf material, and RNA samples representing a total of 100mg of material were pooled together. Total RNA from pycnia and aecia stages were isolated from 100mg of infected larch needles using a protocol adapted to RNA isolation from conifers (Chang et al., 1993). A DNase I (QIAgen) treatment was applied to all samples according to the manufacturer’s recommendations to eliminate traces of genomic DNA. Electrophoretic RNA profiles and RNA quality were assessed with the Experion analyser using the Experion Standard-sens analysis kit (Bio-Rad, Marnes la Coquette, France) before sample shipment and in Beckman’s facilities (Beckman Coulter Genomics, Grenoble, France) prior RNA-sequencing. The following RIN numbers were recorded before Illumina libraries preparation: 8.6, 8.3, 8.8, 6.9, 7, 6.8, 7.6, 8.6 and 8.3, corresponding to replicates R1, R2, R3 of basidia, pycnia and aecia samples, respectively. cDNA synthesis and RNA-Seq library construction were performed by Beckman Coulter Genomics using the Illumina TruSEQ RNA seq kit following the manufacturer’s recommendations. Each library was quantified by qPCR and sequenced with the Illumina HiSeq2500 platform as paired-end 100nt reads. Data acquisition was performed in Beckman Coulter Genomics facilities according to standard Illumina’s procedures. On average, 114 million paired-end reads were produced per sample (ranging from 95 to 129 million reads per sample, Table S1). The complete expression dataset is available at the NCBI Gene Expression Omnibus repository under the series number GSE106863.

### Reference genome and annotation

The *M. larici-populina* isolate 98AG31 genome (version 1.0; US Department of Energy Joint Genome Institute; http://genome.jgi.doe.gov/Mellp1/Mellp1.home.html; Duplessis et al. 2011a) was used as a reference. The RNAseq analysis was performed using the catalog of 16,399 *M. larici-populina* isolate 98AG31 reference transcripts. Automatic annotations such as Gene ontology (GO), Kyoto Eukaryotic Genes and Genomes (KEGG) and euKaryotic Orthologous Group (KOG) annotations were recovered from the reference genome website. The expert manual annotation of specific gene categories (e.g. carbohydrate-active enzymes, CAZymes; lipases; proteases; transporters) performed by the poplar rust genome consortium was used (Duplessis et al. 2011a). The repertoire of SSPs previously described (Hacquard et al. 2012) was extended to include all SPs of unknown function independently of their protein length as described in Lorrain et al. (2015).

### Gene expression analysis

All reads per sample were mapped onto the 16,399 *M. larici-populina* transcripts using the RNAseq analysis tool of CLC Genomics Workbench 6.5.1 (QIAgen) with default parameters, except for the similarity fraction which was set at 90% across 90% of the read length. Reads showed an average per base Phred quality score higher than 30 over their 100 nt length in all samples and no trimming was applied before mapping. The number of unique reads mapped per transcript (counts) was recorded for each sample and the summary of mapping statistics is presented in Table S1. A saturation curve analysis was performed with the rarefy function of the VEGAN package (Oksanen et al. 2017) to determine whether the sequencing depth of our data was sufficient. R packages DESeq2 (v1.10.1) and edgeR (3.12.0) were used to conduct differential expression analysis of *M. larici-populina* transcripts (Love et al., 2014; Robinson et al., 2010). Counts were normalized using the size factor method proposed by Anders and Huber (2010). Differentially expressed genes (DEGs) were then identified using both packages (padj<0.05) and the intercept was considered for further analyses with DESeq2 count values as the reference dataset. The R package WGCNA was used to generate clusters of co-expression (R script available on demand) among significantly DEGs (Langfelder and Horvath, 2008). For this purpose, only DEGs found between all three stages were considered.

### KOG enrichment analysis

KOG enrichment analysis was performed as previously described in Hacquard *et al.* (2013). Secreted proteins of unknown function were extracted from the KOG categories “No Hit” and "Function unknown” to define a new KOG category that we termed “Secreted proteins”. Over-represented KOG categories in basidia, pycnia and aecia up- or down-regulated genes (3-fold change significantly DEGs) were determined as respect to the global gene distribution. Significance of overrepresented KOG categories in each condition was estimated using the Fisher’s Exact Test and False Discovery Rate corrected by Benjamini-Hochberg Test (FDR < 0.05).

### Comparison between *M. larici-populina* larch and poplar infection transcriptomes

Expression data for *M. larici-populina* dormant and germinating urediniospores, and for time-course infection of poplar leaves were recovered from a previous transcriptome analysis performed with whole-genome custom oligoarrays (Duplessis et al., 2011b). Expression data were recovered for 14,883 *M. larici-populina* transcripts showing expression values higher than the background level. The expression sets were reduced for comparison purpose as follows: expression values in dormant and germinating urediniospores were grouped in a unique "Urediniospores” stage (highest expression value of the two stages retained) and expression values during the poplar leaf time-course infection at 24, 48, 96 and 168 hours post-inoculation (hpi) were pooled into a unique “Poplar” stage (highest expression value during the time-course retained). Larch infection expression datasets were similarly reduced into two sets: expression value in basidia was considered as a unique larch-infecting spores set (“Basidia”) and expression values during larch needles infection (i.e. pycnia and aecia) were grouped into a unique “Larch” stage (highest expression value retained). We used a quantile normalization to estimate and compare transcript distributions within the oligoarrays and RNAseq datasets with the R package Limma (Ritchie et al., 2015). Regarding the specific gene families comparison, some gene members or gene families had non-available expression records from oligoarrays. This was due to the oligonucleotide inability to discriminate between transcript species. These genes were thus not included in the comparative expression analysis.

### RTqPCR

A random set of 22 SSP genes was selected to assess RNAseq expression profiles by RT-qPCR. Primer sequences and efficiency are presented in Table S2. Primer design, first-strand cDNA synthesis and PCR amplification were performed as previously described (Duplessis et al., 2011b; Hacquard et al., 2010). Transcript expression levels were normalized with *M. larici-populina* alpha-tubulin and elongation factor reference genes (Hacquard et al., 2011b), considering the specific efficiency of each primer (Pfaffl, 2001). For each gene, in order to compare profiles measured by oligoarrays and RTqPCR, expression ratios were calculated between relative levels in each experimental condition and the average expression level across conditions.

## Results

### *M. larici-populina* transcriptome of larch infection related stages

To gain insights into the genetic program of *M. larici populina* during larch infection, we performed high depth RNAseq at three distinct stages: basidia at 0 dpi, pycnia at 7 dpi and aecia at 14 dpi (Figure 1). An average of 114 million reads per sample and more than 1 billion reads in total were collected (Table S1) and mapped onto 16,399 *M. larici-populina* transcripts. Overall, 16,386 transcripts had at least one read assigned. The proportion of reads mapped ranged from 10.3 to 59.3% in pycnia replicate 2 and basidia replicate 1, respectively (Table S1). This reflected the different proportion of fungal transcripts in plant tissues. Each replicate reached a plateau in rarefaction curves independently of the stage (Figure S2), indicating a sufficient sequencing depth to capture a full transcriptome at each stage. Principal component analysis of DESeq2 rlog-transformed normalized counts showed homogeneous biological replicates (Figure S3A). The first axis (94% of the variance) separated basidia from aecia and pycnia, whereas the second axis (3% of the variance) separated aecia from pycnia. One aecia replicate was apart from the others, probably because of the higher proportion of null counts observed for this replicate and the overall heterogeneity of the biological material at this stage compared to basidia and pycnia. Normalized counts distributed in a similar manner across all the samples (Figure S3B). Average expression values were then considered for the basidia, pycnia and aecia stages. Overall, these data confirmed the good quality of the RNAseq approach for further comparative analysis.

### Gene expression profiles correlate in pycnia and aecia (larch infection stages)

To further explore differences between basidia and larch *in planta* stages, we analysed global gene expression profiles and differentially expressed genes (DEGs). Pycnia and aecia stages occurring in larch needles showed overall closer expression profiles, whereas they are more dispersed when compared with basidia (Figure 2A-C). In total, 10,204 transcripts (62% of the 16,399 predicted *M. larici-populina* transcripts) were commonly detected in basidia, pycnia and aecia (Figure 2D). Only 230, 696 and 48 transcripts were detected exclusively in basidia, pycnia and aecia, respectively. Pairwise comparisons were performed between basidia and larch *in planta* stages, and transcripts with normalized fold change and padj<0.05 in at least one paired comparison were deemed as significantly differentially expressed genes (DEGs; up- or down-regulation). In total, 7,743 transcripts were significantly differentially expressed in at least one stage compared to another (Table S3) and 7,031 and 7,079 transcripts were significantly differentially expressed between pycnia and basidia or aecia and basidia, respectively (Table 1, Table S4). Fewer DEGs (2,141) were detected between aecia and pycnia. We conclude that the closely related profiles recorded in pycnia and aecia support the expression of a stable genetic program *in planta* in larch.

**Figure 2:**
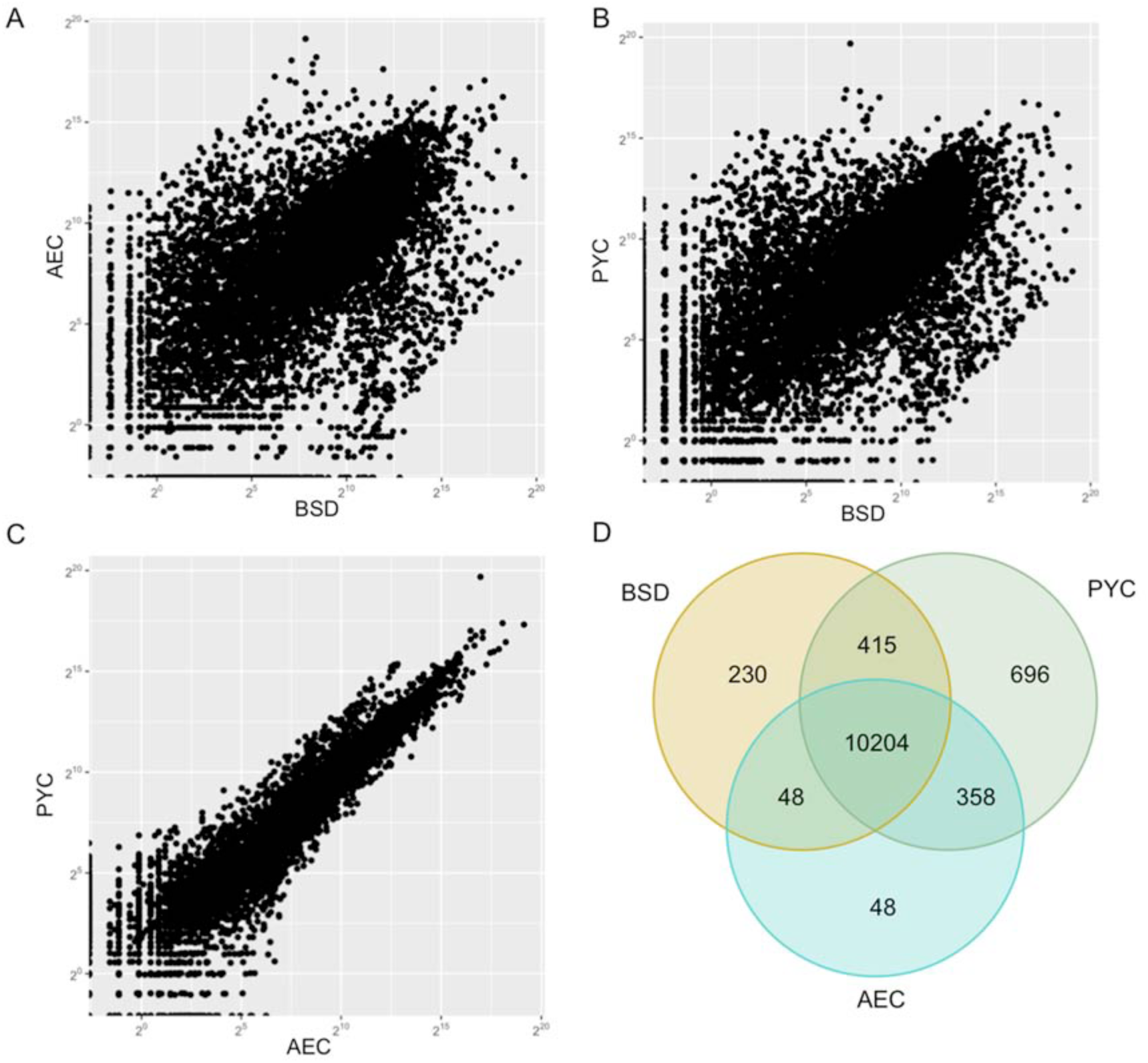
RNA-seq analysis of basidia (BSD), pycnia (PYC) and aecia (AEC) of the rust fungus *Melampsora larici-populina*. (A) Scatter plot of rlog transformed sequencing counts of transcripts expressed in AEC *versus* BSD; (B) in PYC *versus* BSD and (C) in PYC *versus* AEC. (D) Venn diagram showing the number of genes detected by at least one read per sample in each condition Basidia (gold), Pycnia (green) and Aecia (blue).

**Table 1:** Number of *Melampsora larici-populina* genes significantly differentially expressed in basidia (BSD), pycnia (PYC) and aecia (AEC).

| **Regulated genes<sup>a</sup>** | **PYC vs.**<br>**BSD** | **AEC vs.**<br>**BSD** | **AEC vs.**<br>**PYC** |
| --- | --- | --- | --- |
| Genes significantly differentially expressed ($p < 0.05$) | 7031 | 7079 | 2141 |
| Up-regulated | 3698 | 3503 | 1045 |
| > 3-fold | 2406 | 2280 | 90 |
| > 10-fold | 1526 | 1316 | 5 |
| > 100-fold | 522 | 442 | 0 |
| > 1000-fold | 105 | 67 | 0 |
| Down-regulated | 3333 | 3576 | 1096 |
| < 3-fold | 1872 | 2128 | 539 |
| < 10-fold | 849 | 1053 | 265 |
| < 100-fold | 319 | 429 | 27 |
| < 1000-fold | 57 | 126 | 0 |

<sup>a</sup> Down- and up-regulated genes were classified according to their fold-change ratio. Transcripts with a significant $p$ -value ( $< 0.05$ ) and more than a threefold change in transcript level were considered as significantly differentially expressed.

### Secreted proteins is the only overrepresented category among DEGs detected on larch

To discriminate between sets of *M. larci-populina* genes expressed in basidia and during larch infection, we conducted a co-expression network analysis (Langfelder and Horvath, 2008). This correlation-based network analysis identifies clusters of genes with expression profiles that are more linked to each other than they are with genes outside the cluster. Among 1,548 significantly DEGs (padj<0.05), we identified six clusters of transcripts sharing co-expression profiles (Figure 3A, Figure S4). The clusters positively correlated with basidia were negatively correlated with *in planta* stages pycnia and aecia (Figure 3A, Figure S4). Clusters 1 and 2 were significantly positively correlated with basidia and represented 38.8 % of the DEGs (661). The cluster 3 was significantly positively correlated with pycnia, representing 21.3% of the selected DEGs (330). Clusters 4 to 6 were significantly positively co-expressed in aecia with 39,8% of DEGs (617; Figure 3A). In total 61.2% of the dEgs were significantly co-expressed *in planta* compared to basidia. Three over four clusters (clusters 3, 4 and 5) were significantly negatively correlated with basidia, while positively correlated with *in planta* stages (Figures 3A and S4).

**Figure 3:**
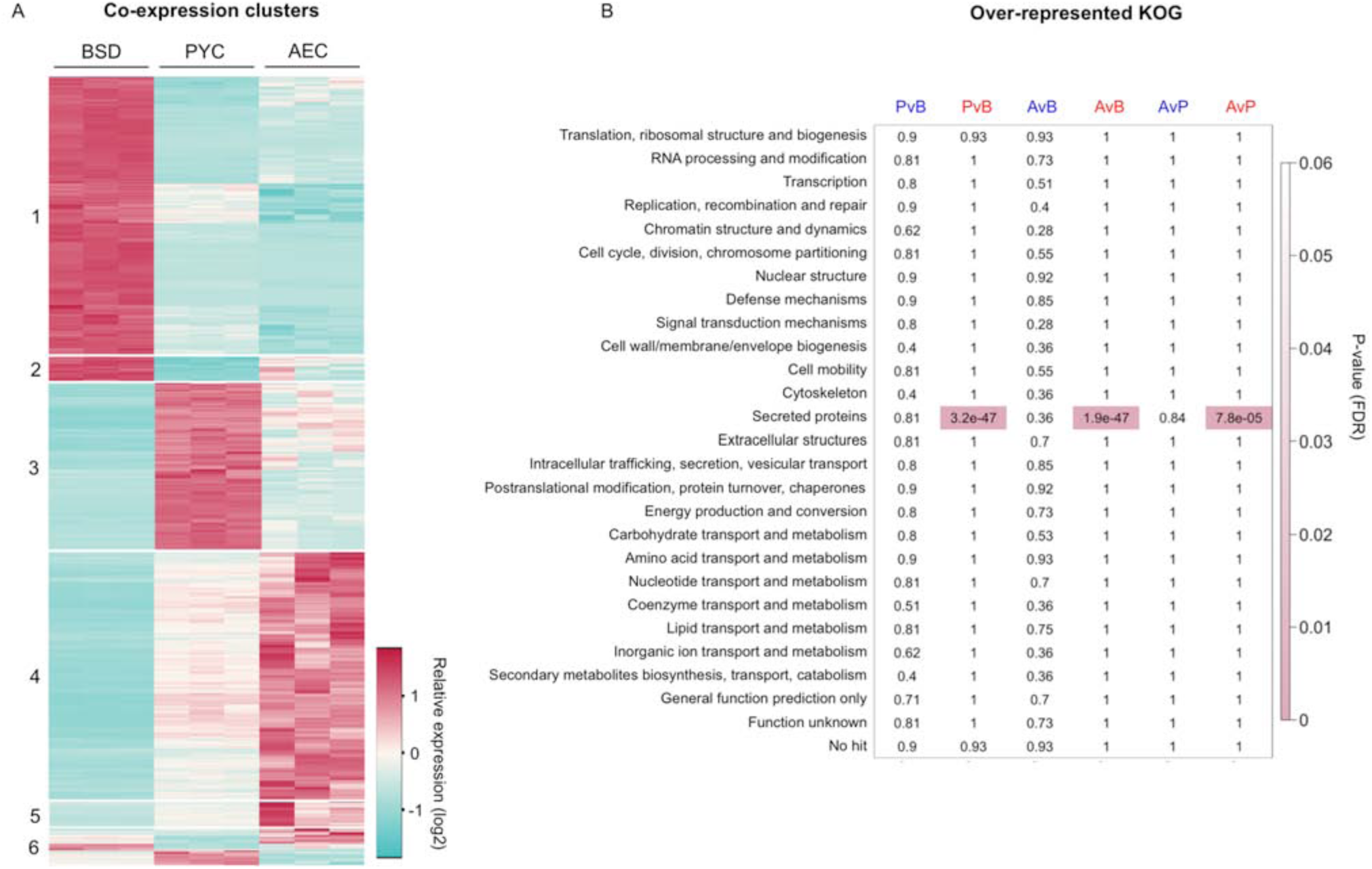
Co-expression analysis and KOG enrichment analysis of differentially expressed *Melampsora larici-populina* genes in basidia and larch infection stages. (A) Transcript profiling of 1,548 *M. larici-populina* significantly differentially expressed genes at the three stages BSD, PYC and AEC (DESeq2; pajd <0.05). Relative expression profiles are represented as log2 of normalized counts across three replicates of BSD, PYC and AEC stages, from over-expression (red) to under-expression (blue). Using co-expression network analysis (WGCNA), genes were grouped into six clusters of co-expression. (B) FDR p-values matrix from over-represented KOG categories of 3 fold-change down-regulated genes (blue) and 3 fold-change down-regulated genes (red). Significantly enriched categories (FDR<0.05) are highlighted in pink.

To further understand the transcriptome reprogramming occurring between basidia and larch *in planta* stages, we conducted an enrichment analysis of KOGs in pairwise comparisons of regulated genes. Transcripts regulated with at least |≥ 3fold-change| were considered (Table 1). We also created a specific KOG-SP category for secreted proteins of unknown function. Among up-regulated genes in aecia versus pycnia (1,045 DEGs), aecia versus basidia (3,503 DEGs) and pycnia versus basidia (3,698 DEGs) pairwise comparisons, the only significantly enriched category is KOG-Sp (Figure 3B). SPs represented 28.9%, 22.4% and 22.1% of aecia versus pycnia, aecia versus basidia and pycnia versus basidia up-regulated genes, respectively. SP encoding genes were over-represented in up-regulated genes of aecia and pycnia stages compared with basidia. We also performed RT-qPCR on 22 randomly selected SP genes to confirm their RNAseq expression profiles (Figure S5). A good correlation could be observed between RNAseq and RT-qPCR for pycnia and aecia whereas differences are noticeable for basidia, which may relate to the intrinsic nature of the biological material (dead poplar leaves). Our results show that more SP genes are expressed during larch *in planta* stages compared to basidia.

### Comparison of *M. larici-populina* transcriptomes on larch and poplar highlights subsets of specically and commonly expressed genes

To identify *M. larici-populina* genes specific to larch or poplar infection we compared transcriptome profiling of larch stages to previous expression studies performed on the telial host (Duplessis et al. 2011b). We simplified our dataset to four explicit stages: basidia as a larch-infecting spore stage, LARCH as the *in planta* stage on the aecial host, Urediniospores as a poplar-infecting spore stage and POPLAR as the *in planta* stage on the telial host. Expression data recorded for poplar-related stages were obtained with custom oligoarrays. Thus, direct calculation of fold-change was not possible between larch- and poplar-related stages due to the different technologies used. However, the transcriptomes were complete on both hosts (Duplessis et al. 2011b and Figure S2), allowing direct comparison to determine the presence of specifically expressed genes (at least count of 1 by RNAseq or above background value on oligoarrays). Overall, 14,883 transcripts were detected in poplar- or larch-related stages, and 8,035 transcripts are expressed in all four fungal stages (Figure 4A). A total of 1,436 transcripts were only detected on basidia and LARCH, and 1,531 transcripts were detected on Urediniospores and POPLAR. Interestingly, 389 genes were specifically detected in LARCH and POPLAR, revealing a set of *in* planta-specific genes. In total, 2,967 (20%) genes are specifically expressed on larch or poplar *in planta* stages.

**Figure 4:**
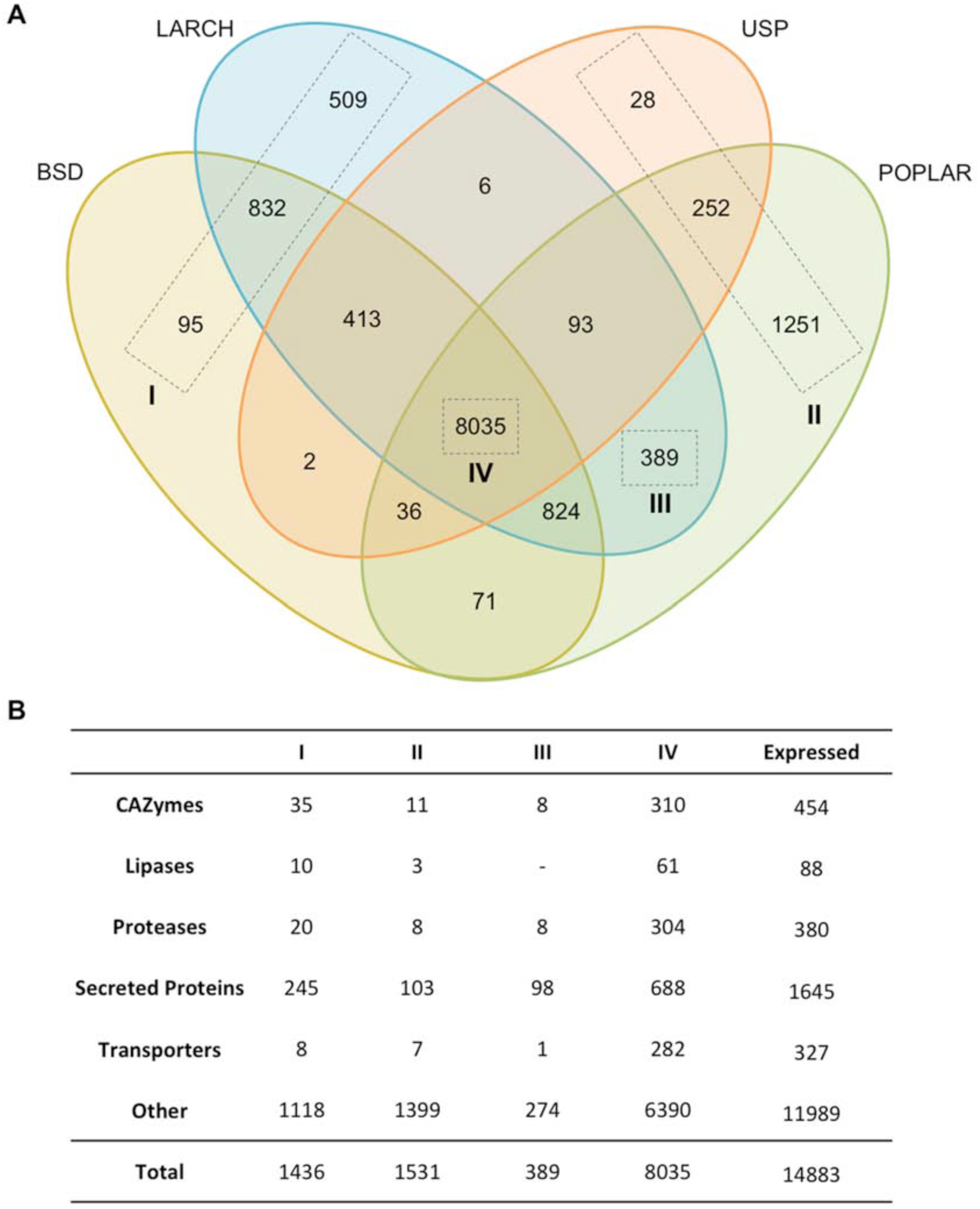
Comparative transcriptomic analysis between four stages of *Melampsora larici-populina* life cycle. (A) Venn diagram showing the number of genes expressed in BSD (basidia), LARCH (pycnia and aecia, i.e. during larch needle infection), USP (dormant and germinating urediniospores) and POPLAR (during poplar leaf infection). Group I refers to genes specifically expressed in larch-related stages; group II refers to genes specifically expressed in poplar-related stages; group III refers to the genes specifically expressed *in planta* and group IV refers to genes detected in all four stages. (B) Number of genes expressed in selected annotated gene categories related to host infection: carbohydrate-active enzymes (CAZymes), lipases, proteases, secreted proteins and transporters in selected groups I, II, III and IV. The total numbers of expressed genes for each selected annotated category is shown on the right. Total numbers of genes belonging to selected groups I, II, III and IV are shown below the table.

### Detailed analysis of gene categories expressed on larch and poplar reveals SP genes specifically expressed on each hosts

To determine whether *M. larici-populina* uses specific pathways or functions during colonization of larch and poplar, we surveyed expression profiles of gene categories. Firstly, no major difference could be determined on one host or another regarding the expression of KOG cellular categories (Figure S6). Manual exploration of KEGG annotations revealed a similar result for metabolic pathways (Table S3). Secondly, of 6,660 genes of unknown function (excluding SP genes addressed below) with expression information, 58% were expressed at all stages, whereas 13% (863) and 14.5% (967) presented a specific expression pattern on larch and poplar, respectively (Figure S7). We scrutinized cellular categories previously annotated in the *M. larici-populina* genome (Duplessis et al. 2011a). Among those, 282 transporters, 310 CAZymes and 304 proteases were expressed at all four stages. Few genes in these surveyed categories were specifically detected in larch- and poplar-related stages. Likewise, only 8 transporters were detected only in Basidia and LARCH, including three auxin efflux carriers and one oligopeptide transporter, which belong to expanded rust fungal gene families (Duplessis et al. 2011a; Figure 4B; Table S3). A total of 35 CAZyme genes were detected in Basidia and LARCH stages, including 22 glycoside hydrolases; and 20 proteases were detected at these stages, including 8 peptidase inhibitors. Even fewer genes were specifically detected in USP and POPLAR, with 7, 11 and 8 transporters, CAZymes and proteases, respectively. Overall, less than 10% of CAZymes, proteases, lipases or transporters are specifically expressed in polar or larch stages. Among mating-related genes, pheromone receptor genes were expressed in both hosts and one gene showed a very high expression in LARCH. Four pheromone precursor genes showed a very high expression in LARCH compared to other stages and were in the top 1% most highly expressed genes in aecia (Table S3), whereas mating compatibility is established between pycniospores and receptive hyphae. In total, 60% of the specifically expressed genes on poplar and on larch are of unknown function.

Since SP genes were significantly over-represented in *M. larici-populina* genes expressed on larch (this study) and on poplar (Duplessis et al. 2011b), we particularly considered their expression profiles during larch and poplar infection. Among 1,645 expressed SP genes, 688 (42%) were detected at all developmental stages (Figure 4B). Transcripts specifically detected either on larch or poplar contained an important proportion of SP-encoding genes, accounting for 17 and 25% of the total number of genes specifically expressed at these stages, respectively. A lower proportion of SP genes were specifically detected on the telial host poplar, compared to the aecial host larch. Even if the proportion of SP genes expressed at all stages was important (688; 42% of 1,645 SP genes), it represented only 8.6% of the 8,035 transcripts expressed at all four stages. Moreover, of the top 10% genes preferentially highly expressed on larch or on poplar *in planta*, 18% are secreted proteins and among them, 14% are SSPs (Figure 5). In comparison, SPs and SSPs represent only 4 and 5% rescpectively of other expressed genes. Altogether, these results indicate an enrichement of SSPs among genes highly expressed during poplar or larch infection.

**Figure 5:**
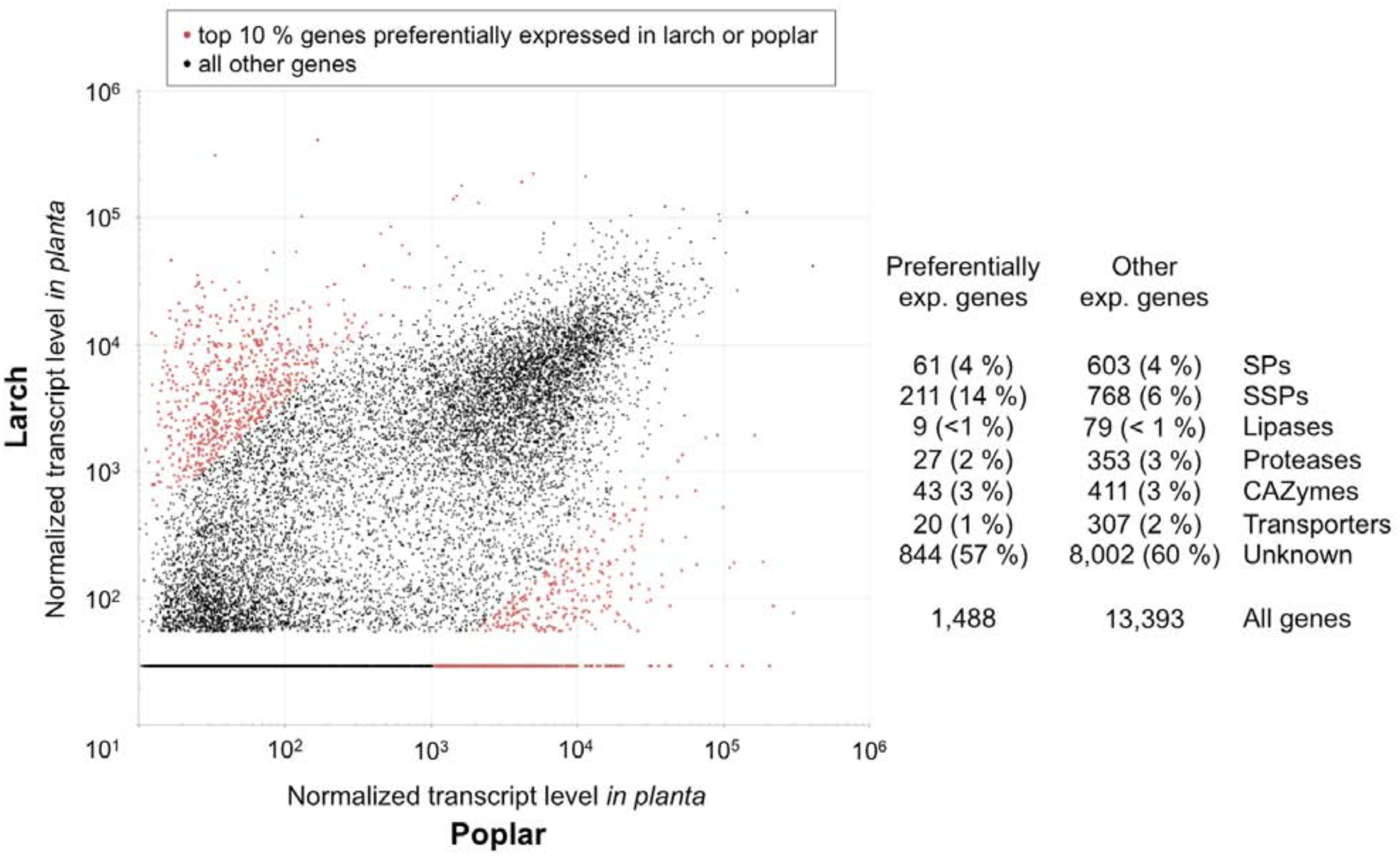
*Melampsora larici-populina* genes preferentially expressed *in planta* on larch and on poplar. Scatter plot representing the normalized gene expression level of 14,883 *M. larici-populina* genes expressed *in planta* on poplar (x-axis) and on larch (y-axis). The red dots represent the top 10% genes preferentially expressed on poplar (5%) and on larch (5%). Numbers and percentage of genes in secreted proteins (SPs), small secreted proteins (SSPs), lipases, proteases, CAZymes, transporters and genes of unkown function (Unknown) of the top 10% genes and of other expressed genes are detailled. The original expression data for rust-infected poplar leaves and rust-infected larch needles were quantile normalized to allow comparison.

### Gene members of *M. larici-populina* SSP families present specific patterns of expression on larch and on poplar

We took the opportunity of the detailed analysis of SSP gene families in *M. larici-populina* (Duplessis et al., 2011a; Hacquard et al., 2012) to further explore their specific expression patterns on the two host plants. RNAseq and oligoarray expression data were quantile-normalized and compared to determine whether entire families or members of families are host-specific (Figure 6). Overall, 205 and 351 SSP family genes were preferentially expressed *in planta* on larch and on poplar, respectively when the general expression profile of all SSP genes was considered (Figure 6A). When focusing on singletons or families of SSPs grouping two, three or more members, three-fourths of the genes were detected on the two hosts (Figure 6B). The remaining fourth corresponded to genes expressed specifically in one or the other host (Basidia and LARCH or Urediniospore and POPLAR). We further considered large SSP families to identify family-specific expression patterns. Among 64 families of at least four members, 17 (26%) showed a preferential expression on one given host (9 and 8 SSP families preferentially expressed in larch or poplar *in planta* stages, respectively). For instance, the SSP family 1 (111 members) presented 24 members expressed specifically on Basidia and LARCH and 4 only on Urediniospore and POPLAR, the remaining members showing a higher expression in Basidia and LARCH (Figure 6C). On the contrary, the SSP family 9 (11 members) showed a preferential expression on poplar, and only two members presented noticeable expression level on the other host. The SSP family 7 presented a striking profile with almost a half of the gene members expressed in the aecial host larch, whereas the other half was expressed in the telial host poplar. Some of these members presented a sequence identity of 75% and showed opposite expression profiles on the two host plants (e.g. sSp genes 86274 and 91075 expressed in larch and in poplar, respectively; data not shown). In smaller SSP gene families, the different situations exemplified in the Figure 6C were also observed (Table S3). Expression profiles of *M. larici-populina* SSP gene families proved to be diverse with a 60% of genes expressed on the two host plants, 31 % showing preferential expression on one host or the other and 9% of divided families (6 SSP families of more than three members).

**Figure 6:**
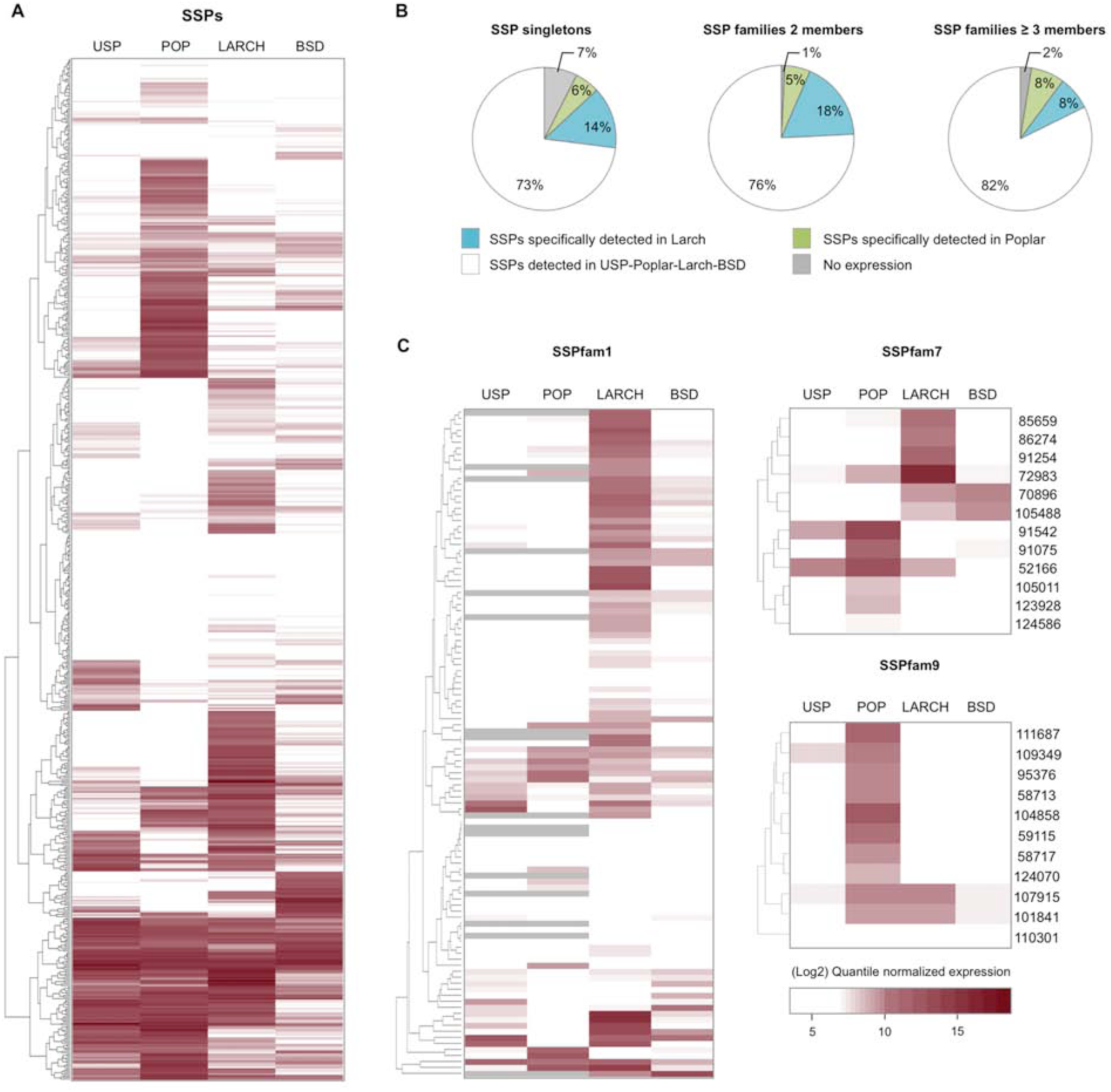
Transcript profiling of *Melampsora larici-populina* SSP genes on aecial and telial host. (A) Heatmap of expression profiles of *M. larici-populina* SSP transcripts. Expression levels (white to red) are shown as quantile normalized expression values (white corresponding to the lowest expression and dark red corresponding to the greatest expression). (B) Diagrams presenting the expression patterns for SSP singletons, SSP gene families with 2 members and SSP gene families with 3 or more members detected in larch- and poplar-related stages (i.e. BSD and LARCH; and USP and POP, respectively). (C) Examples of expression profiles for selected SSP families: SSP family 1, preferentially expressed in larch; SSP family 7, expressed both in poplar and larch; SSP family 9, preferentially expressed in poplar. JGI protein ID numbers are indicated on the right of profiles for SSP families 7 and 9. Grey on the SSPfam1 expression profile indicates non-available information (lack of specific oligomers on the oligoarray).

## Discussion

In the present study, we report on gene expression profiling at three biological stages that are related to infection of larch, the aecial host of the poplar rust fungus *M. larici-populina*, taking advantage of an experimental set-up in semi-controlled laboratory conditions (Pernaci et al., 2014). A full transcriptome was recovered at each of the stages through deep RNA-Sequencing allowing comparisons between stages. In total, 73% of *M. larici-populina* genes were expressed at basidia, pycnia and aecia stages. This is similar to previous findings (76%) where stages related to infection of poplar, the telial host of the fungus were considered (Duplessis et al., 2011b; Hacquard et al., 2010, 2012). Altogether, 62% of the genes were commonly expressed in basidia, pycnia and aecia, however the general profile was closer between the two larch-infection stages pycnia and aecia than with basidia. This suggests an important reprogramming of the fungal transcriptome after the transition to host infection. Although pycnia and aecia stages occur before and after fertilization, major *in planta* structures such as protoaecia are formed before the fertilization between pycniospores and receptive hyphae of compatible mating types (Harder, 1984; Voegele et al., 2009), which may explain the similarity of the general expression profile at these stages. It is worth noticing that mating-related genes were among the most highly expressed genes detected in pycnia. The analysis of the wheat leaf rust genome recently resolved the mating system and loci for a rust fungus (Cuomo et al., 2017). Exploring expression of mating genes on the host on which sexual reproduction occurs would allow a better understanding of the underlying genetic processes.

Heteroecious rust fungi exhibit the most complex life cycle known in fungi. They involve infection of two taxonomically unrelated host plants and production of five different types of spores (Aime et al., 2017). Transcriptomics of rust has been widely applied to asexual stages and spores responsible for epidemics and damages in plantations (Duplessis et al. 2014). There is still scarce information about expression along the whole life cycle of rust fungi, and only a few studies looked at transcript profiles on alternate host plants. In *Puccinia triticina*, EST sequencing and RNA-Seq identified sets of genes expressed on the two host plants, meadow rue (*Thalictrum* spp.) and wheat, however at a limited scale (Xu et al., 2011; Cuomo et al., 2017). RNA-Seq was applied at a higher depth to *Cronartium ribicola* spores collected from the two host plants, *Pinus monticola* (aeciospores) and *Ribes nigrum* (urediniospores), and compared to two infection stages in western white pine (Liu et al., 2015). Although no reference genome is available for *C. ribicola*, the RNA-Seq approach could identify a large proportion of *C. ribicola* genes expressed in all situations (61%) with only a few (4-5%) specific to each separate stages (Liu et al., 2015).

In our study, the significant over-representation of SPs among regulated genes between stages was striking and mirrors previous findings on the poplar host (Duplessis et al., 2011b; Hacquard et al., 2010, 2012). This shows that the expression of SP genes is important for larch infection and central to biotrophy on both hosts. Many effectors of fungal pathogens are secreted proteins, including avirulence effectors identified in the flax rust fungus *Melampsora lini* (Tyler and Rouxel, 2011; Sharpee and Dean, 2016, Duplessis et al., 2012). Rust fungi exhibit large secretomes composed of expanded gene families and SSP genes specifically expressed *in planta* represent priority candidate effectors (Aime et al., 2017; Petre et al., 2014). Previous expression profiling reports of poplar infection by *M. larici-populina* have identified long list of putative effectors that are key candidates for functional analysis (Hacquard et al., 2010; Joly et al., 2010; Duplessis et al. 2011b; Hacquard et al. 2012; Petre et al., 2012; Hacquard et al., 2013). No expression data was recorded before during infection of larch by *M. larici-populina.* Whether specific sets of effectors may be required for rust fungi to infect their different hosts was so far unaddressed (Shulze-Lefert and Panstruga, 2011). Only a fraction of SSP genes was specifically expressed on poplar or on larch, in a similar fashion for singletons or multigene families and a larger portion is expressed on the two hosts. Our results confirm and largely extend the observations made in *P. triticina* or *C. ribicola* regarding specific gene expression (Xu et al., 2011; Liu et al., 2015; Cuomo et al., 2017). Considering that rust effectors are likely to reside among these genes, our results could indicate that a considerable part of *M. larici-populina* effectors may share similar functions on the two hosts.

Some *M. larici-populina* SSP families showed a preferential expression on a single host, and more striking was the profile of families exhibiting members whose expression was host-specific. Families showing host-specific expression could reflect a diversification under the pressure of resistance genes or the evolution of a new effector function targeting a particular process not represented in the alternate host. Regarding gene families combining specificity of expression towards the two hosts, such as the *M. larici-populina* SSP7 family, we may observe adaptation of subsets of effectors i) with different functions in the cells of the two hosts, or ii) with a conserved function but different targets in the two plants, or iii) having the same target that evolved differently in the two plant lineages from a unique ancestral target. These results open many avenues for speculation about the evolution of SSP and candidate effectors families in rust fungi, but it remains to determine whether these particular families correspond to *bona fide* effectors.

Beyond candidate effectors, other SP genes such as the CAZymes showed specific expression in poplar or in larch. Similar observations were made for *C. ribicola* infecting *Ribes* spp. or pine (Liu et al., 2015). It is known that plant-cell wall varies in terms of composition, architecture and integrity across plant species (Kubicek et al., 2014). CAZymes are essential enzymes for decomposing polysaccharides from plant cell walls and leaf or needle cuticles (Henrissat et al., 2017). Rust fungal hyphae needs to penetrate through the plant cuticle in the aecial host and to go through the plant cell wall to form haustoria within the host cell cavity in both hosts (Ragazzi et al., 2005). Different small sets of CAZymes were expressed in poplar and larch, which could reflect the adaptation to the host polysaccharide composition in a conifer and a dicot. On the contrary, the survey of central metabolic pathways or cellular categories did not reveal major differences. *M. larici-populina* seems to express a conserved genetic program during the biotrophic growth in larch and poplar tissues suggesting a common strategy for the acquisition and utilisation of nutrients derived from the two hosts. A surprising feature of rust fungal genomes is the proportion of genes of unknown function with no homology out of the Pucciniales (Duplessis et al. 2011a, Aime et al. 2017). Among 6,660 *M. larici-populina* genes of unknown function with expression record, more than 10% are specifically expressed on poplar or on larch, whereas 58% are expressed on both. A better dissection of infection on the two host plants could help to draw hypotheses about processes unknown yet in which these genes could be involved, e.g. early colonization, biotrophic growth or sporulation. Moreover, systematic comparative transcriptomics in different heteroecious rust fungi may provide new leads to unravel new functions related to biotrophy.

When transcriptome is considered across the whole *M. larici-populina* life cycle, we could detect expression for 86% of the genes and a total of 2,047 genes were never detected. Although this relatively small number of genes may contain annotation errors, it is likely that some may be specific of sections of the rust biological life cycle not explored in this or in previous studies. Telia are overwintering structures adapted to drastic environmental conditions in which important biological processes such as karyogamy and meiosis occur (Mendgen, 1984). A small proportion of genes specifically expressed in telia were previously reported in *M. larici-populina* (Hacquard et al. 2013), which could be explained by the fact that transcriptome profiling was carried out in early differentiating telia. This telia stage was not included for comparison in the present study because it was obtained from environmental samples different from the reference strain 98AG31 (Hacquard et al. 2013). When compared with the present study, only 117 genes could be detected as telia-specific (data not shown). More specific profiles can be expected at later stages of telia differentiation when karyogamy and meiosis occur or during entry and exit of winter dormancy. Moreover, the infection of the aecial host is far more complex than in the telial host with the establishment of distinct fungal cell types in a coordinated manner (Harder, 1984; Littlefield and Heath, 1979). Only three specific stages were selected for this work and specific expression of other genes may be expected during the course of the interaction with the aecial host, as previously illustrated in poplar (Duplessis et al., 2011b; Hacquard et al., 2010, 2012).

Other fungal biotroph possess the capacity to interact with multiple host species with various levels of specificity. Mycorrhizal fungi are biotrophs that engage in mutualistic associations with their host plants by forming specific infection structures (van der Heijden et al. 2015). During the colonization process, they deliver symbiosis effectors to establish long lasting beneficial interactions (Martin et al., 2016). Large-scale transcriptome studies have shown that ectomycorrhizal fungi also present specific expression of large sets of SSP genes of unknown function (Kohler et al. 2015). Similar to our observations with *M. larici-populina*, mycorrhizal fungi express common and specific sets of SSP genes during their interaction with different host plants. This has been shown for ectomycorrhizal fungi in different interactions between *Suillus* spp. and *Pinus* spp. (Liao et al., 2016) and *Laccaria bicolor* with poplar and Douglas fir (Plett et al., 2015). The same was observed for the arbuscular mycorrhizal fungi *Rhizophagus irregularis* and *Gigaspora rosea* colonizing different host plants (Kamel et al., 2017). The fungal root endophyte *Piriformospora indica* establishes different colonization strategies in a host-dependant manner, i.e. biotrophy in *Arabidopsis* and saprotrophy in barley (Lahrmann et al. 2013). Transcriptome analysis of root colonization in these plants shows the expression of common and specific sets of SSP genes. Although the identified SSPs are different in these different fungal species, there is a common trend for their expression during infection, particularly during the establishment of biotrophic interactions, suggesting that they may participate in defining the level of specificity with different host plants.

## Acknowledgements

The authors would like to acknowledge the ‘Investissements d’Avenir’ program (ANR-11-LABX-0002-01, Lab of Excellence ARBRE) and the Joint Genome Institute (Office of Science of the U.S. Department of Energy under contract no. DE-Ac02-05cH11231) for the sequencing and annotation of the genome of the poplar rust fungus *Melampsora larici-populina* and access made through the mycocosm portal. Emmanuelle Morin and Annegret Kohler at INRA Nancy are acknowledged for their bioinformatic support. Cecile Lorrain is supported by a young scientist grant (CJS) from INRA.

## Author contributions

SD, AH and PF planned and designed the research; CD, SH and JP performed experiments; CL, CM, BP and SD analysed data; CL, CM, BP and SD wrote the manuscript; all authors read and approved the final manuscript.

---

**Figure S1.**
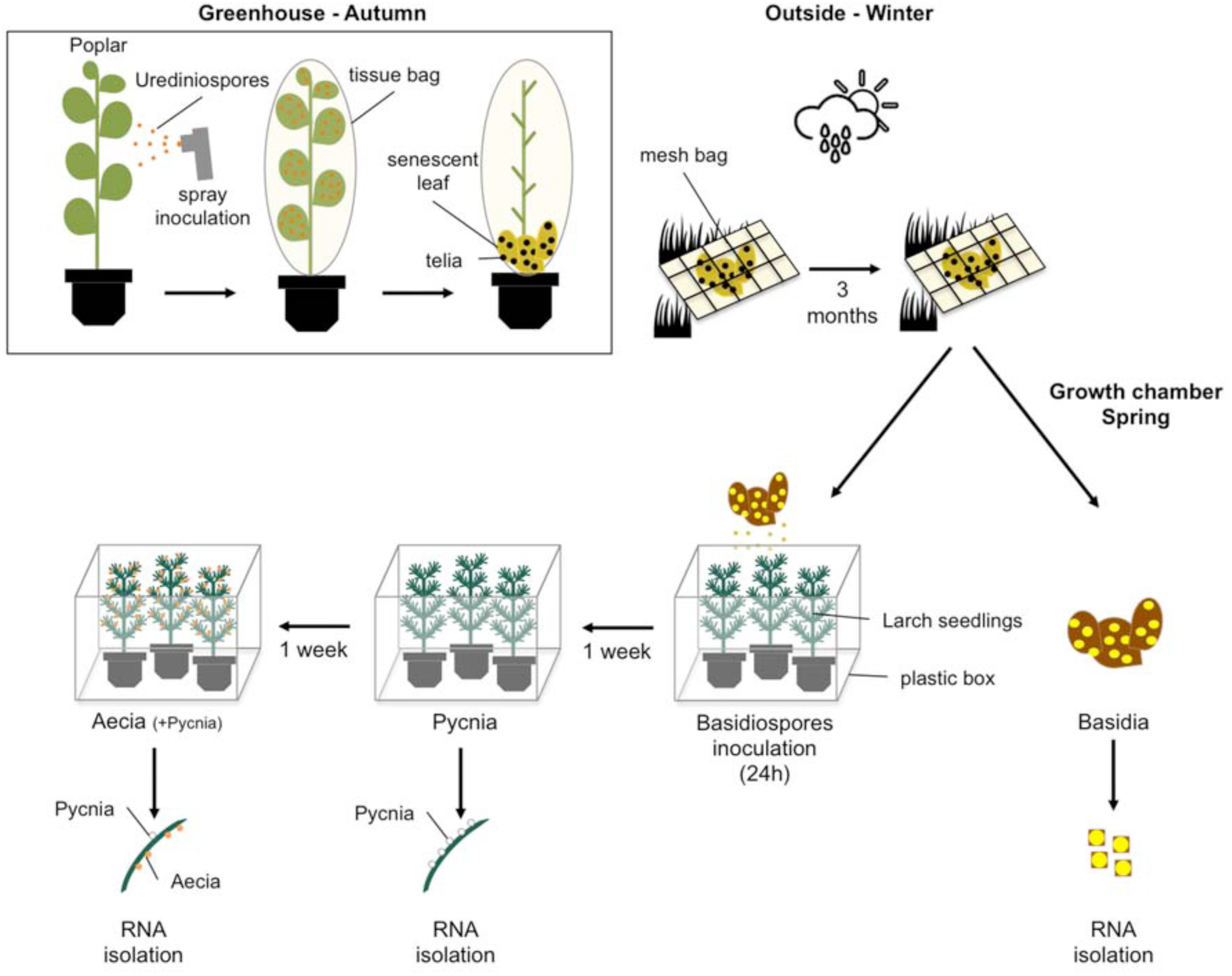
Experimental procedure for basidia production, inoculation of larch seedlings and collection of pycnia and aecia in *Melampsora larici-populina* for RNA isolation according to Pernaci et al. 2014. Leaves of young poplar ‘Robusta’ cv. plants grown in greenhouses were inoculated with urediniospores of *M. larici-populina* isolate 98AG31 in September. After dehiscence, senescent infected leaves were let stand in tissue bags until telia developed. Leaves were then collected and placed outside in mesh bags to let teliospores overwinter under natural conditions. Leaves were used the following spring as primary source of basidiospore inoculum to infect young larch seedlings. From April to May, dead poplar leaves showing large black crust area corresponding to telia were placed in Petri dishes on moist paper at room temperature until yellow basidiospores can be seen at the surface of the leaves. A mixture of overwintered telia remains and freshly produced basidia and basidiospores were collected and snap frozen in liquid nitrogen. This corresponds to the basidia stage. *Larix decidua* seedlings were grown for three months in a greenhouse. Basidiospore-producing poplar leaves were placed over the larch seedlings in plastic boxes kept in the lab at room temperature in order to ensure infection of young soft needles. After one day, poplar leaves were removed to avoid continuous release of basidiospores over time. Larch needles showing high density of nectar droplets were collected and snap frozen in liquid nitrogen one week after inoculation. These samples contain different types of haploid fungal cells such as *in planta* infection hyphae and pycniospores (i.e. flexuous hyphae and pycniospores) and correspond to the pycnia stage. Boxes containing larch needles were kept closed most of the time and small flies (dark winged fungus gnats, Sciaridae, Diptera) naturally present in the substratum of the larch seedlings ensured fertilization in nectar droplets. Aecia and aeciospores were visible two weeks after inoculation on larch needles on the opposite side of pycnia. Needles showing a large number of aecia were collected and snap-frozen in liquid nitrogen (stage Aecia). A few nectar droplets were still visible on some needles at this stage.

**Figure S2:**
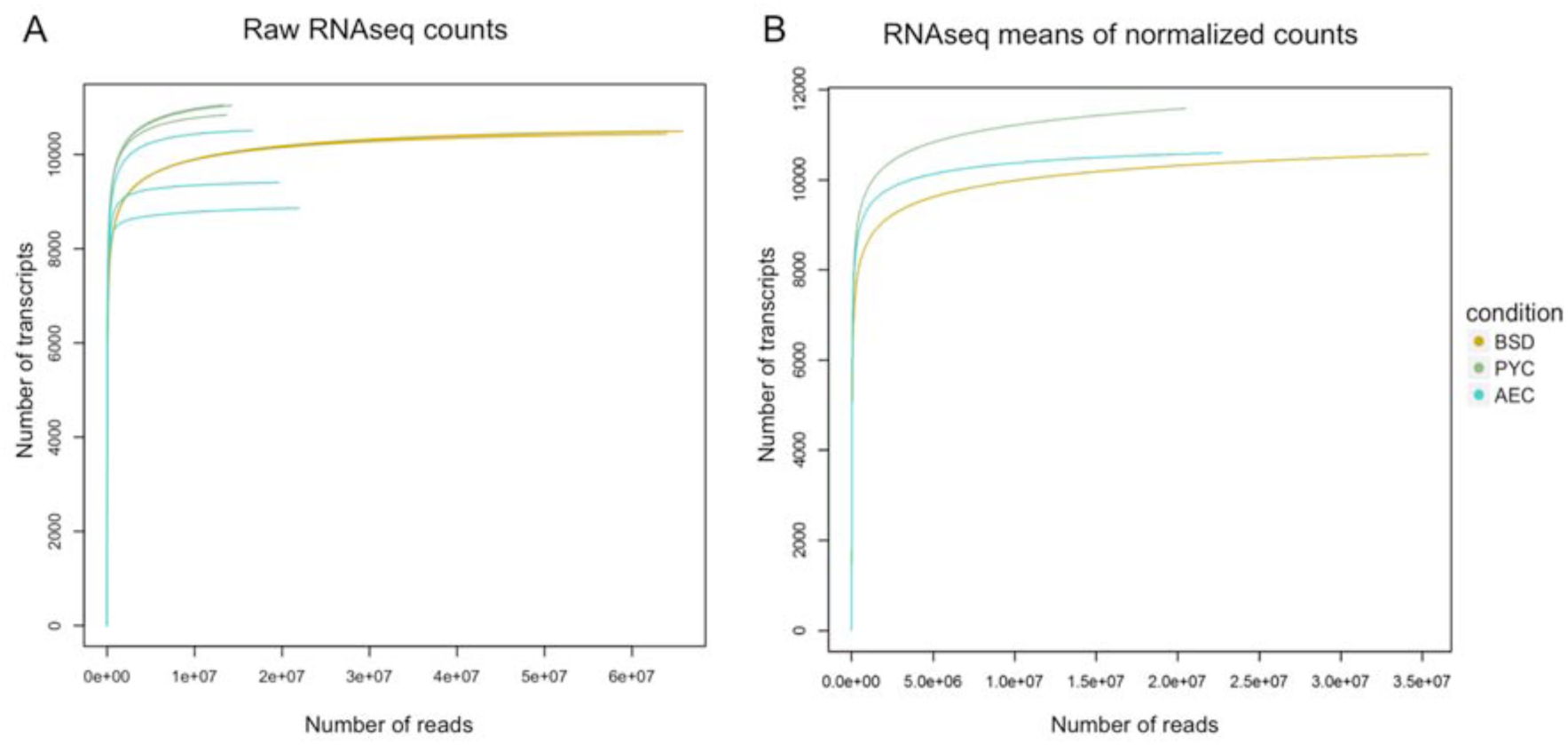
Sequencing saturation curves showing high depth RNA sequencing of *Melampsora larici-populina* basidia, pycnia and aecia stages. The number of sequencing reads (x-axis) attributed to a given number of transcripts randomly selected from the total pool of sequencing reads (y-axis) is shown for the raw RNAseq counts of the three biological replicates of BSD, PYC and AEC stages (A) and for the means of the normalized counts of the three biological replicates (B).

**Figure S3:**
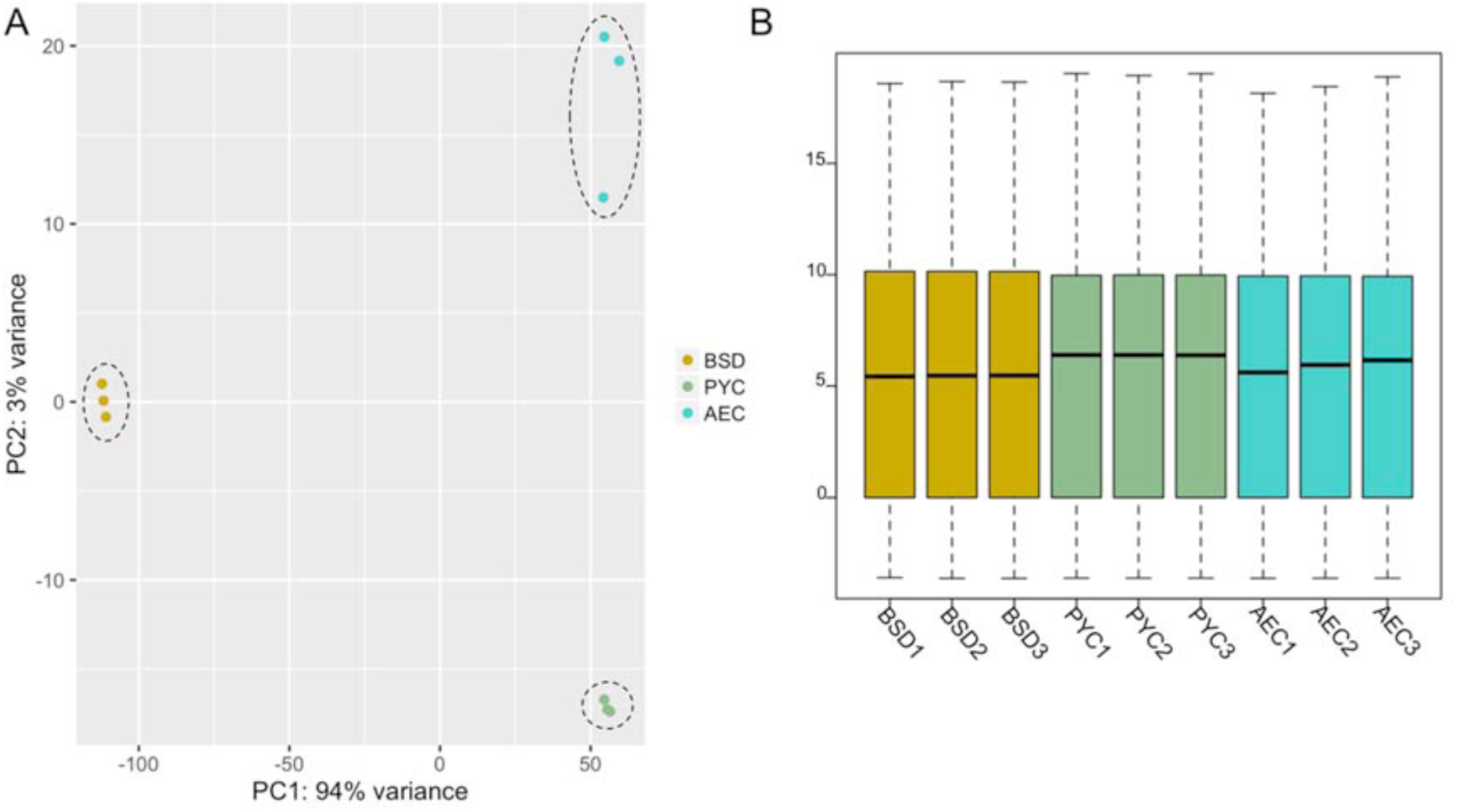
Comparison of RNAseq libraries between *Melampsora larici-populina* basidia, pycnia and aecia. (A) Principal component analysis (PCA) of *M. larici-populina* transcripts levels measured in BSD (gold), PYC (green) and AEC (blue) stages using RNA-sequencing. Reads detected per transcript (counts) were normalized using the size factor method proposed by Anders and Huber (2010) and rlog transformed before performing the PCA. The PCA plot places biological replicates along the two PC axes explaining 94% (x-axis) and 3% (y-axis) of the variance within samples. (B) Boxplots showing the distribution of sequencing counts for three biological replicates of basidia (gold), pycnia (green) and aecia (blue) stages. The top and bottom of boxes correspond to the 25 and 75 % quartiles, respectively. The middle line represents the median (50% quartile). Bottom whisker refers to the 1.5 interquartiles range of the lower quartile and top whisker indicates the 1.5 interquartile range of the upper quartile.

**Figure S4:**
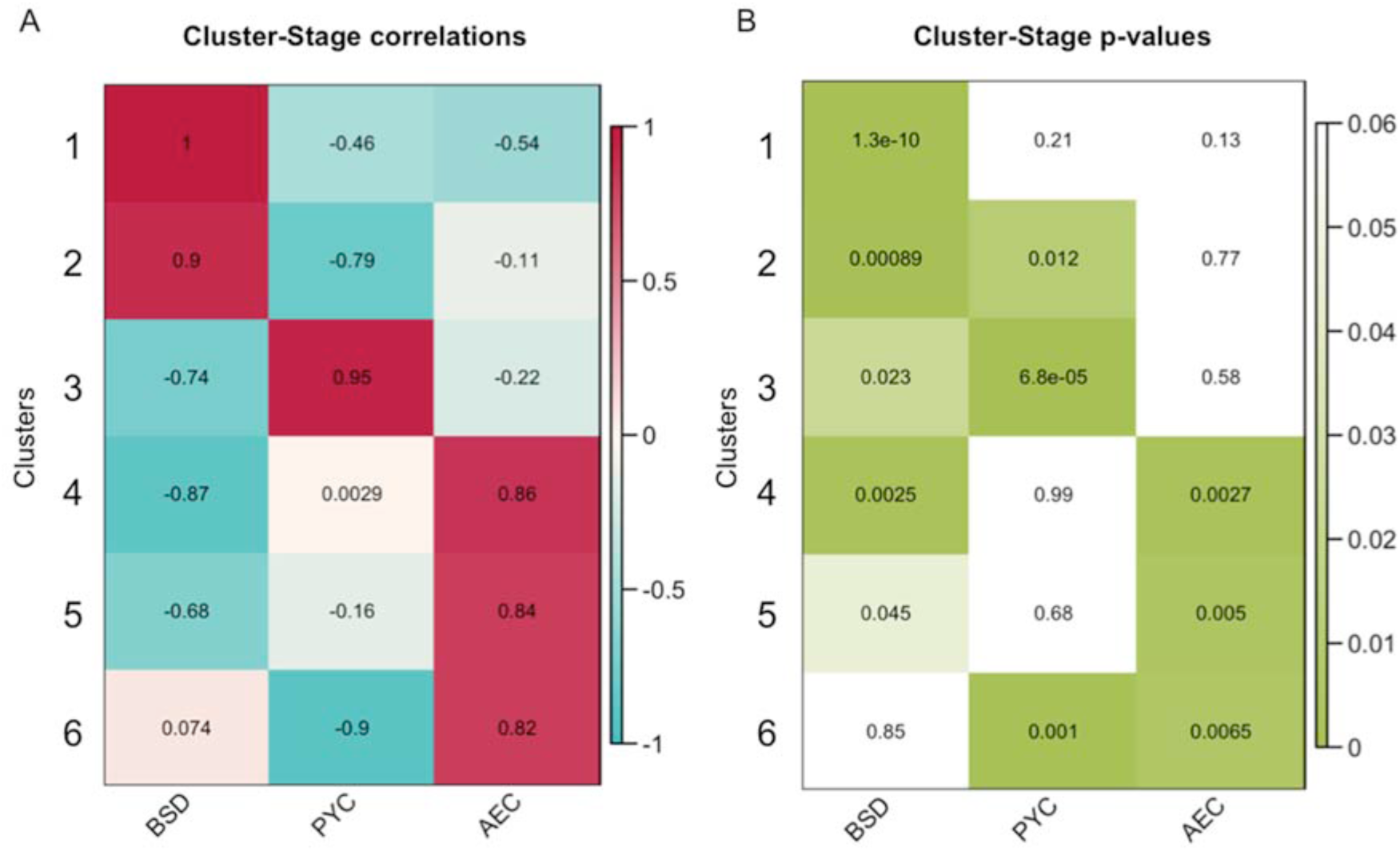
Expression patterns of six *Melampsora larici-populina* clusters of regulated genes. (A) Each row in the matrix corresponds to a cluster of co-expressed genes in each condition (BSD, PYC and AEC). Clusters are correlated from -1 (negative correlation in blue) to 1 (positive correlation in red). (B) *p*-values matrix of cluster-stage correlations with significant correlations (*p*-value<0.05) highlighted in green.

**Figure S5:**
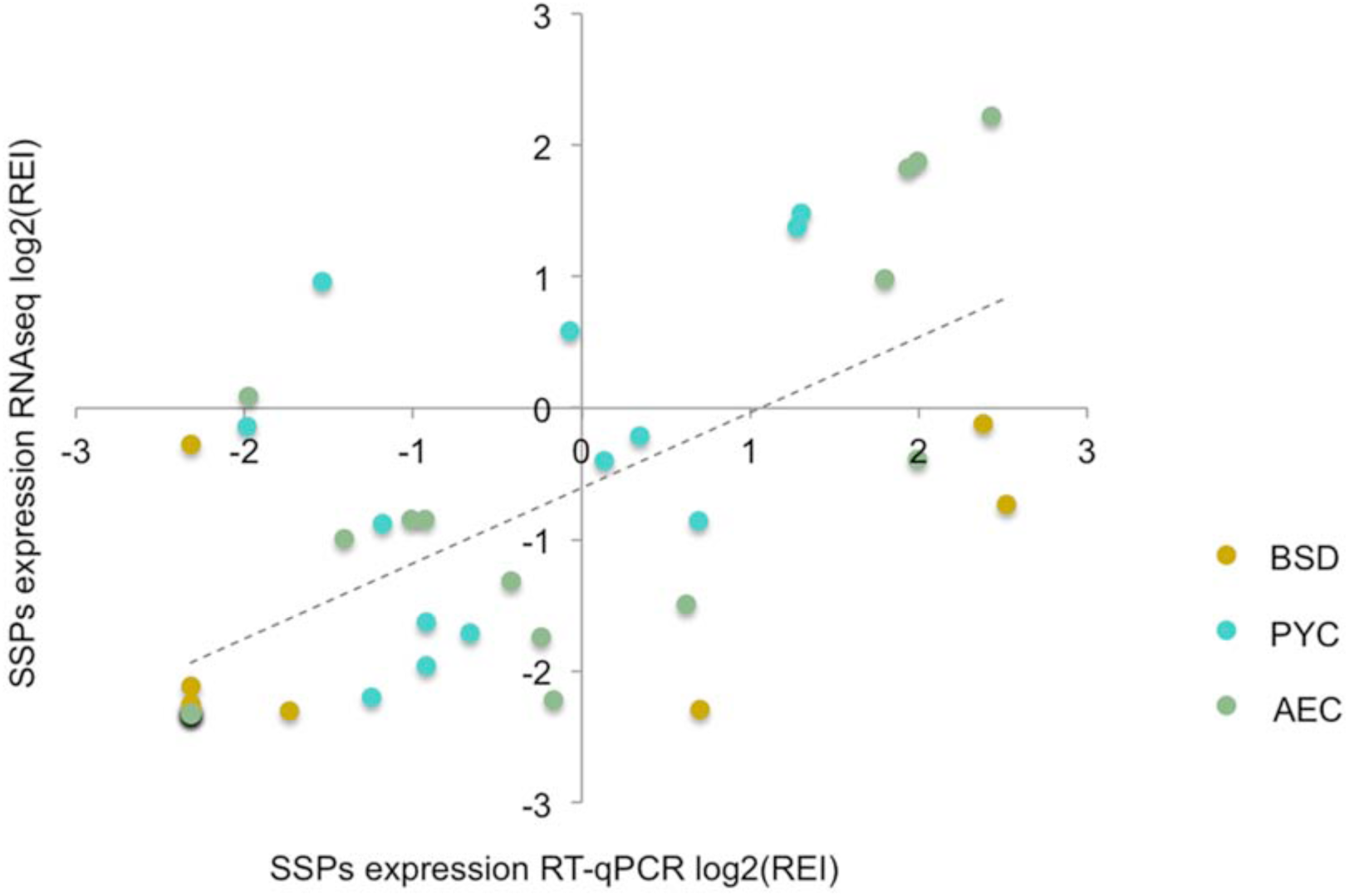
Comparison of SSP expression profiles monitored by RT-qPCR and RNAseq in basidia, pycnia and aecia. For each SSP, RT-qPCR expression levels were obtained from biological triplicates and normalized with *Mlp-a-TUB* and *Mlp-ELF1a* reference genes. For each gene, in order to compare profiles measured by oligoarrays and RTqPCR, expression ratios were calculated between the relative expression level in each experimental condition and the average expression level across all conditions. Ratios were Log2-transformed for presentation.

**Figure S6:**
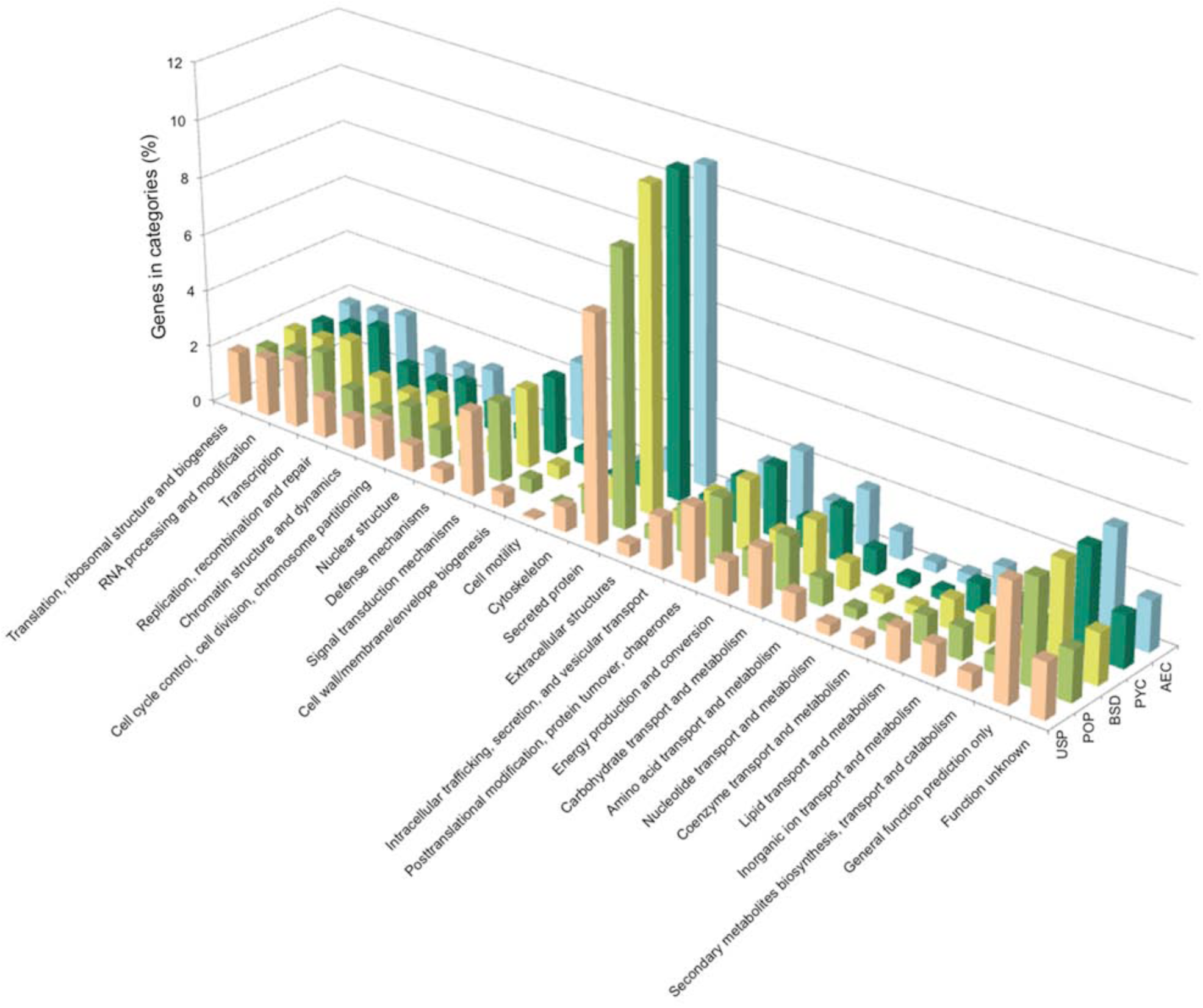
KOG categories distribution among expressed genes at different *Melampsora larici-populina* life cycle stages. Distribution of KOG categories among expressed genes detected at different stages of the *M. larici-populina* life cycle: USP, urediniospores; POP, infected poplar leaves; BSD, basidia; PYC, pycnia; AEC, aecia. The KOG category “No Hit” was the most abundant for all stages and was not included on graph for a better data representation. Note the inclusion of a category for “secreted protein”. Distribution on the y-axis is provided as percentage values of the total number of genes detected at each stage. The lower proportion in the “secreted protein” category for USP and POP compared to other stages is due to non-available data on oligoarrays (i.e. gene families with no specific oligomers for given members).

**Figure S7:**
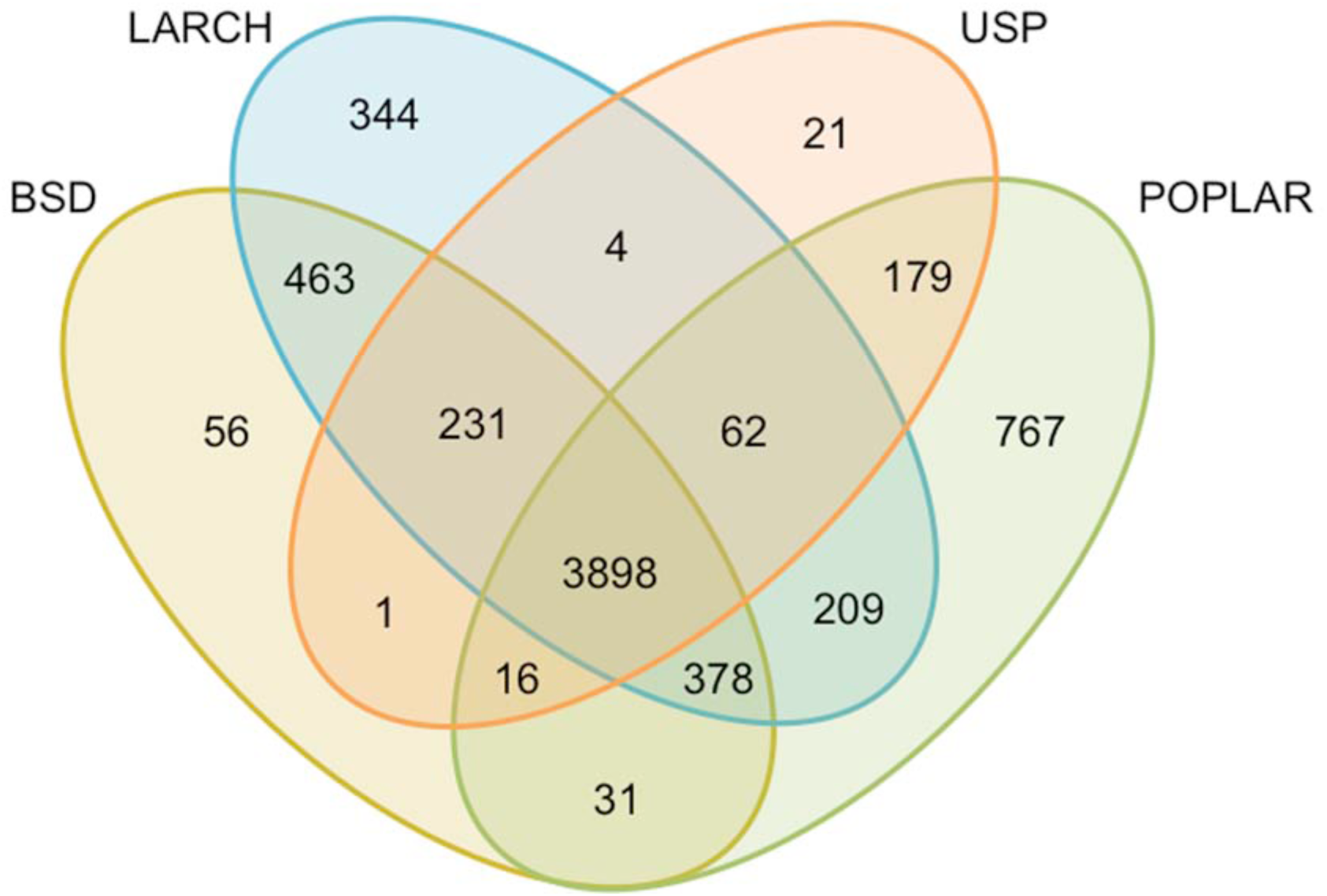
Comparison of expression profiles of genes of unknown function across *Melampsora larici-populina* life cycle. Venn diagram showing the number of genes of unknown function expressed in stages BSD (basidia), LARCH (pycnia and aecia, i.e. during larch needle infection), USP (dormant and germinating urediniospores) and POPLAR (during poplar leaf infection).

**Table S1:**
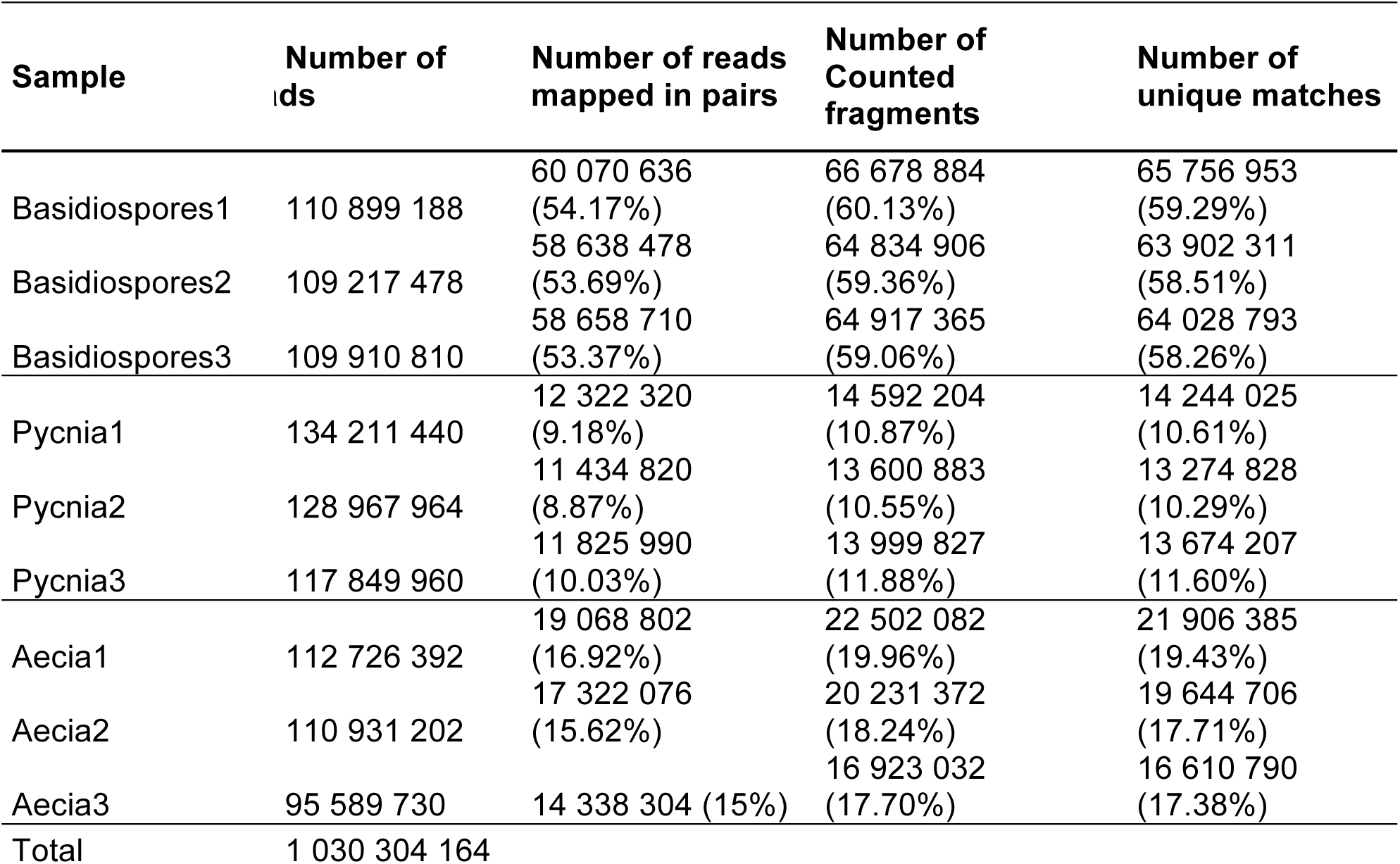
General mapping information of RNA-sequencing data against *Melampsora larici-populina* genes. Illumina reads of the three biological replicates of basidia, pycnia and aecia samples were compared to the catalogue of predicted *M. larici-populina* transcripts. The initial number of reads, the number of reads mapped in pair, the total number of counted fragments and the number of unique matches are shown. The total number of reads across all nine data points is also indicated.

**Table S2:**
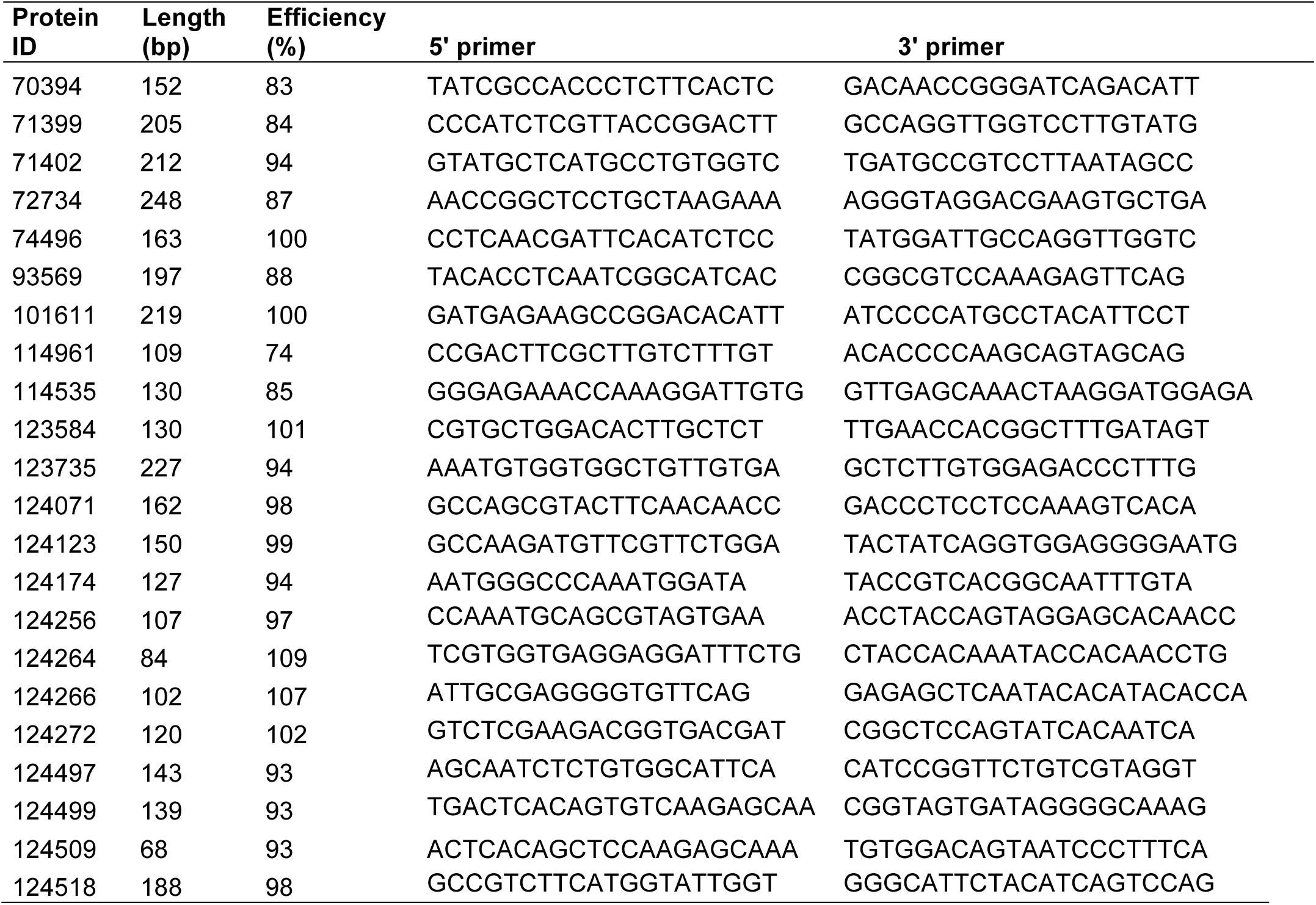
RT-qPCR primers designed to determine expression profiles *of Melampsora larici-populina* SSP genes. Specific 5′ and 3′ primers for 22 *M. larici-populina* SSP genes are detailed, along with the amplicon length (bp) and primers efficiency (%).

**Table S3: *Melampsora larici-populina* transcripts annotation and expression information.**

For each of the 16,399 *M. larici-populina* transcripts, the following annotation and expression information are provided: JGI ProteinID, protein size (amino acids), manually annotated categories (SP/SSP, secreted proteins/small secreted proteins; CAZyme, carbohydrate active enzymes; CytP450, cytochrome-P450), category information (details about subfamilies within a given category according to specific databases, i.e. CAZY for cazymes, MEROPS for proteases and TransportDB for transporters), SSP families with numbers of members (according to Hacquard et al. (2012)), gene ontology (GO) annotation, Kyoto encyclopedia of genes and genomes (KEGG) annotation, eukaryotic orthologous groups annotation (KOG; ID, description and class), expression in urediniospores (USP), poplar, basidia (BSD), pycnia (PYC) and aecia (AEC) (expression in USP and poplar corresponds to the highest expression level in dormant and germinated urediniospores and in a time-course infection of Beaupré leaves, respectively, according to Duplessis et al. (2011b) and larch corresponds to the highest expression level in pycnia and aecia), quantile-normalized expression values for USP, poplar, BSD and larch stages (USP_norm, POPLAR_norm, BSD_norm and LARCH_norm, respectively). In columns for USP and poplar, the red color indicates expression value below background. Automatic annotations for GO, KEGG and KOG databases were retrieved from the *M. larici-populina* genome webportal (https://genome.jgi.doe.gov/Mellp1/Mellp1.home.html) and manual annotations were defined according to Duplessis et al. 2011a.

**TableS4: *Melampsora larici-populina* differentially expressed genes between the three larch infection-related stages.**

The table presents the fold change expression levels between the three larch-infection stages basidia, (BSD), pycnia (PYC) and aecia (AEC) for genes found significantly differentially expressed between at least two stages with DESeq2 (adjusted *p*-value padj <0.1). JGI *M. larici-populina* proteinID, log2FoldChange, FoldChange and padj are detailed.

